# Sustainable Technology for the Fabrication of Liposomal Phases

**DOI:** 10.64898/2026.05.09.724055

**Authors:** Ayshwarya Ravikumar, Swathi Shanmugam, Anirban Polley

## Abstract

Liposomes are self-assembled lipid vesicles capable of encapsulating both hydrophilic and hydrophobic therapeutics, making them versatile platforms in drug delivery and biomedical technology. In this study, the limitations of the classical thin-film hydration method were critically evaluated, and a sustainable, systematically optimized strategy was established for generating defined liposomal lamellar phases. Hydration conditions were optimized, and 4 mL of buffer per 10 mg of lipid was determined to be optimal for effective rehydration and improved statistical reliability of vesicle measurements. A refined probe-sonication protocol (20% amplitude, 5 s ON/55 s OFF pulse) enabled controlled transformation of multivesicular vesicles into stable multilamellar and unilamellar vesicles at net ON-times of 90 s and 185 s, respectively, without overheating or contamination. In addition, a Python-based machine-learning tool was developed for vesicle size characterization. Collectively, these optimizations provided a reproducible and sustainable framework for preparing liposomes across different lamellar phases.

## 1. Introduction

A liposome is an artificially synthesized, self-assembled lipid vesicle composed of a single bilayer or multiple concentric bilayers enclosing an aqueous core. Alec Bangham first introduced the field of liposomology and characterized liposome structure in the mid-1960s. Since then, liposomes have been extensively studied as one of the simplest biomimetic systems, resembling miniature cells without the complexities of nuclei or cytoplasm (Sessa and Weissmann, 1968; Bangham et al., 1974; Adler and Schiemann, 1985; Bibi *et al*., 2011; Aranda-Lara *et al*., 2020; Andra *et al*., 2022).

The size of liposomes ranges from ∼20 nm to several µm, while the lipid bilayer thickness is typically 3–5 nm (Liu et al., 2022). Phosphatidylcholine (PC) lipids constitute ∼40–50% of cell membranes (van der Veen et al., 2017). In this study, dimyristoylphosphatidylcholine (DMPC), a commonly used lipid with a low main phase transition temperature (Tc ≈ 23 °C), is employed for liposome preparation (Drabik et al., 2020). Based on lamellarity, liposomes are classified as unilamellar vesicles (ULVs), multilamellar vesicles (MLVs), and multivesicular vesicles (MVVs) (Giuliano et al., 2021). ULVs may be further categorized as SUVs (<100 nm), LUVs (100–1000 nm), and GUVs (>1 µm), although such size distinctions are not the focus of the present study (Nsairat et al., 2023).

Liposomes are widely explored in drug delivery as they encapsulate hydrophilic drugs within the aqueous core and hydrophobic drugs within the lipid bilayer, thereby enhancing stability, biocompatibility, and membrane penetration (Gomez and Hosseinidoust, 2020; Umbarkar et al., 2021; Mehta et al., 2023). They interact with cells through endocytosis, exocytosis, and lipid exchange, and serve as vaccine adjuvants and targeted nanocarriers for proteins, nucleic acids, imaging agents, and cancer therapeutics (Wang et al., 2019; Zhang and Sun, 2021; Fulton and Najahi-Missaoui, 2023).

Various macroscale methods—such as thin-film hydration, extrusion, reverse evaporation, ethanol injection, electroformation, freeze-drying, and double emulsion— are widely used for liposome preparation (Šturm and PoklarUlrih, 2021). However, these approaches provide limited control over vesicle size and morphology (Danaei et al., 2018), whereas biological systems require highly defined geometries for specialized function (Choi et al., 2023). Moreover, precise control over cargo release remains a critical challenge, as systemic use of liposomes is often limited by rapid clearance, instability, and unintended drug release (Nagayasu et al., 1999; Kim and Jeong, 2021).

The article is structured as follows. First, the methodology employed for liposome preparation is described in detail. Next, the main results are presented, including the influence of uniform lipid thin-film formation on vesicle morphology, the optimization of rehydration buffer volume, and the calibration of pulsed probe sonication for the formation of distinct liposomal phases. Finally, the findings are summarized, and a concluding perspective is provided outlining key challenges and potential future directions for advancing liposome-based systems.

## 2. Methods

### 2.1 Liposome Preparation by Thin-Film Hydration

Different methods can be employed to prepare various liposomal phases. In the present study, the thin-film hydration technique (Bangham method) was employed for lipid vesicle preparation. Dimyristoylphosphatidylcholine (DMPC) was selected due to its low main phase transition temperature (T_c_ = 23 °C) (Drabik et al., 2020).

Lipids were dissolved in chloroform, followed by solvent evaporation to form a thin lipid film. The dried film was hydrated with HEPES buffer (Lu and Qi, 2021; Lombardo and Kiselev, 2022) and mechanically agitated to generate multivesicular vesicles (MVVs). Subsequent probe sonication was performed to downsize MVVs into multilamellar vesicles (MLVs) and unilamellar vesicles (ULVs).

Vesicle morphology and phase identification were carried out using bright-field microscopy (BF), confocal microscopy (CM), and scanning electron microscopy (SEM) (Fig. S1 in S1 of Supplementary Information (SI)) (Robson *et al*., 2018). These imaging techniques enabled quantification of vesicle number, size distribution, lamellarity, and structural characteristics (Lujan *et al*., 2019).

### 2.2 Preparation of Lipid Thin Film: Influence of Film Uniformity on Vesicle Morphology

Thin lipid films were prepared by either air drying or rotary evaporation to evaluate their effectiveness in producing uniform films.

For film preparation, 10 mg of DMPC was dissolved in 1 mL chloroform to ensure complete solubilization. When fluorescence microscopy was performed, 18:1 Liss Rhod PE was added at a concentration of 1 µM during this step (Shohda et al., 2015). Other lipid-compatible fluorescent dyes were selected based on spectral compatibility and lipid composition. Glassware cleaning procedures and detailed air-drying protocols are provided in S2 and S3 of SI, respectively (Has and Sunthar, 2020).

#### 2.2.1 Air-Drying Method

Thin films were formed by evaporating chloroform under ambient conditions in different containers to evaluate the effect of geometry on film uniformity and vesicle morphology. Air drying in an Eppendorf tube led to uneven solvent evaporation due to its cylindrical shape, sharp edges, and limited surface area, causing lipid accumulation at the base and formation of thicker regions or sediment-like aggregates, which produced heterogeneous vesicle size distribution upon hydration (Fig. S2(a) in SI). In contrast, a concave watch glass provided a larger evaporation surface and yielded comparatively more uniform vesicle distribution after hydration, despite slightly thicker lipid deposition toward the center (Fig. S2(b) in SI). Overall, container geometry influenced evaporation rate, film thickness, and lipid distribution, and air drying frequently resulted in incomplete solvent removal and film heterogeneity (Hadian *et al*., 2014).

#### 2.2.2 Rotary Evaporation Method

For rotary evaporation, the lipid–chloroform solution was transferred to a round-bottom flask and evaporated under reduced pressure at temperatures above T_c_ using a two-step protocol. Initially, solvent removal was performed at 41 °C and 480 mbar with rotation at 60–70 rpm for 15 min to form a thin lipid film. Subsequently, the temperature was lowered to 36 °C and the pressure reduced to 56 mbar while maintaining 70 rpm for an additional 10 min to enhance drying under higher vacuum (Zhu et al., 2013). Alternatively, residual chloroform was eliminated by brief vacuum desiccation (650 mmHg for 1 min) followed by overnight vacuum. Controlled rotation ensured uniform lipid spreading along the flask surface, and reduced pressure promoted efficient solvent removal. Upon rehydration, these films produced vesicles with homogeneous spatial distribution and narrow size dispersion (Fig. S2(c)), outperforming air-dried films (Fig. S2(a), (b)) by yielding thinner, more uniform films with minimal residual solvent contamination, thereby providing superior control over film quality and vesicle homogeneity.

## 3. Results and Discussion

### 3.1. Optimization of Rehydration Buffer Volume

The lipid films prepared by air drying or rotary evaporation were hydrated in aqueous buffer, such as phosphate-buffered saline or HEPES buffer (Lu and Qi, 2021; Lombardo and Kiselev, 2022). In this study, HEPES buffer, a widely used biological buffer that closely mimics physiological conditions (Naddaf Dezfuli et al., 2014), was employed (composition detailed in S4, SI). After rehydration, samples were incubated at 37 °C for 1 h and vortexed for 1 min prior to bright-field imaging. Direct visualization with controlled dilution was adopted to avoid spatial heterogeneity associated with manual smearing and to ensure reproducible vesicle distribution.

To calibrate vesicle density and minimize overlap, the rehydration volume was systematically varied (0.5, 2, 3, 3.5, 4, and 5 mL for 10 mg lipid). Vesicle areas were extracted from microscopy images (Fig. 1(a)), and the probability distribution, P(A), was determined (Fig. 1(b)), approximating vesicles as circular (A = πr^2^). A cutoff area (A_c_ = 50 µm^2^; r_c_ ≈ 4 µm) was used to distinguish overlapping (A > A_c_) from non-overlapping (A ≤ Ac) vesicles. The overlap fraction decreased with increasing volume and plateaued at 4 mL (Fig. 1(c)), whereas vesicle counts initially increased (2 mL) due to reduced overlap and subsequently declined at higher volumes because of dilution (Fig. 1(d)). At 5 mL, vesicle density became insufficient for reliable imaging. Accordingly, 4 mL was identified as the optimal rehydration volume, balancing high vesicle count with minimal overlap for robust bright-field visualization and quantitative analysis (Zhu et al., 2013).

**Figure 1:**
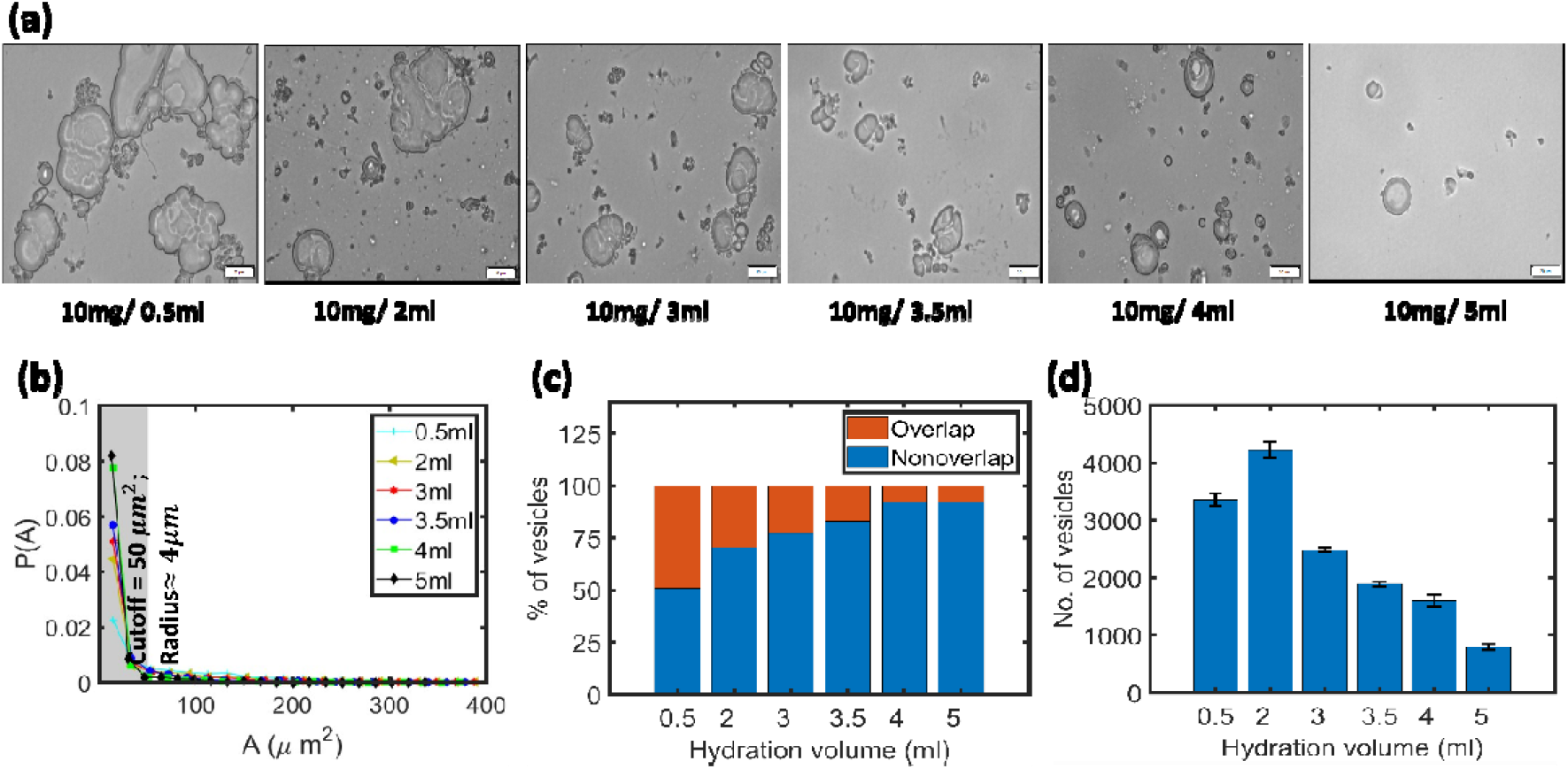
After rehydration of a 10 mg DMPC thin film with buffer (0.5–5 mL), samples were incubated at 37 °C for 1 h, vortexed for 1 min, and imaged by bright-field microscopy. (a) Vesicle distributions at different buffer volumes. (b) Probability distribution of vesicle area, P(A). A cutoff area (A_c_ = 50 µm^2^) distinguishes overlapped (A > A_c_) from non-overlapped (A < A_c_) vesicles (grey). (c) Percentage of overlapped and non-overlapped vesicles versus buffer volume, showing reduced overlap with increasing volume and saturation at 4 mL. (d) Total vesicle count as a function of buffer volume. Error bars denote standard deviation from 20 images.

### 3.2. Optimization of Probe Sonication to Obtain Different liposomal Phases

After rehydration and vortexing (1 min), MVVs were formed (Fig. 1(a); Fig. S4(a) in S7, SI). Vesicle downsizing was performed to obtain MLVs and ULVs. Among available methods—homogenization, sonication, and extrusion (Lombardo and Kiselev, 2022)— sonication is preferred for efficient size control via acoustic disruption of lipid aggregates (Maulucci et al., 2005). Bath sonication (90 min; S5, SI) predominantly yielded MLVs with residual MVVs (Fig. S3, S5, SI), consistent with limitations in power and frequency control (de Freitas et al., 2019; Morrissey, 2001).

Probe sonication (130 W, 20 kHz, 6 mm probe) was therefore employed. Continuous sonication at 30% amplitude for 30 s (Sharma et al., 2023; Zeng et al., 2023) induced heating and metallic contamination (Fig. S4(b) in S7, SI). Ice-bath cooling (Lombardo and Kiselev, 2022) risked phase perturbation due to thermal cycling. To minimize thermal and contamination effects, pulsed protocols were optimized. At 30% amplitude (10 s ON / 50 s OFF), ULVs appeared at net ON-times ≥60 s and were evident at 90 s (Fig. 2(a); Fig. S6 in S9, SI), though heating remained significant.

**Figure 2:**
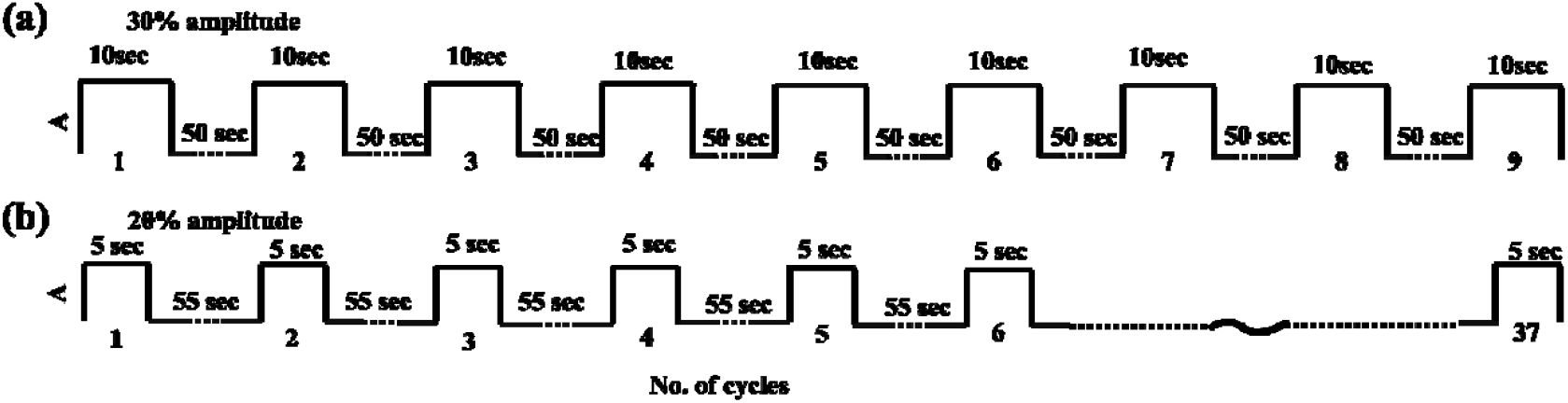
Schematic representation of the pulsed probe sonication, showing amplitude, A with number of cycles. (a) shows the pulsed probe sonication carried out for total 9 cycles (10 second ON and 50 second OFF per cycle) for 30% amplitude, while (b) the same for total 37 cycles (5 second ON and 55 second OFF per cycle) for 20% amplitude.

**Figure 3:**
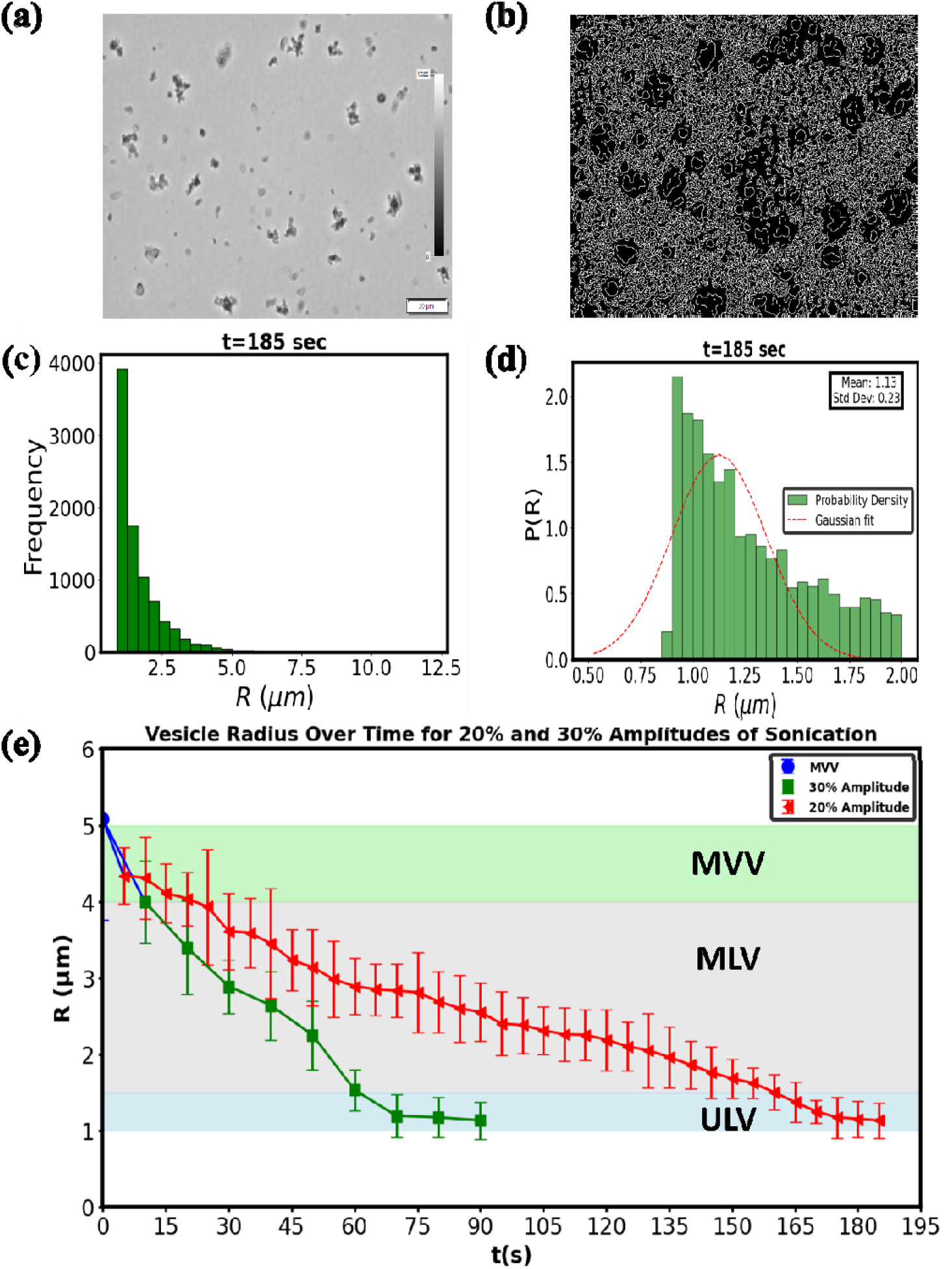
Characterization of vesicles. (a) Bright-field image at an ON-time of 185 s (37th cycle) during pulsed probe sonication at 20% amplitude. (b) Vesicle identification using a custom Python-based machine learning workflow (contour detection). (c) Histogram of vesicle radii (R). (d) Normalized probability distribution, P(R), fitted with a Gaussian function (excluding rare long-tail events) to obtain the mean radius with error. Approximately 20 images (∼4000 vesicles) were analyzed per time point. (e) Evolution of mean vesicle radius with ON-time at 30% and 20% amplitudes, highlighting MVV (green), MLV (gray), and ULV (blue) phases.

A lower amplitude (20%) with 5 s ON / 55 s OFF was then systematically varied (5–185 s net ON-time; Fig. 2(b)). MLVs emerged at 90 s (18th cycle) and ULVs at 160–185 s (32nd–37th cycles). The extended OFF period limited heating and probe erosion. Approximately 20 images per cycle were analyzed using a custom Python workflow (S6; Fig. S7 in S9, SI). Vesicle area (A) and radius (R) were extracted assuming circular geometry (A = πR^2^; 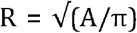.Gaussian fits to P(R) excluding rare long-tail events yielded mean radii with error (Fig. 5c–d). The evolution of mean radius with ON-time (Fig. 5e) classified phases as MVV (R > 4 µm), MLV (R ≈ 1.5–4 µm), and ULV (R < 1.5 µm). Probability distributions (Fig. S5 in S8, SI) confirmed MVV > MLV > ULV, consistent with representative images (Fig. S4 in S7, SI).

Confocal microscopy (Fig. 4(a–d); S1.2 in SI) revealed lamellar structure at 90 s (MLV phase), while SEM at 185 s (Fig. 4(e–f); S1.3 in SI) showed spherical ULVs with slight membrane undulation. Overall, sonication amplitude and duty cycle govern vesicle morphology: higher amplitude accelerates ULV formation but increases thermal risk, whereas low-amplitude pulsed operation enables controlled, contamination-minimized production of distinct liposomal phases (S10 in SI).

**Figure 4:**
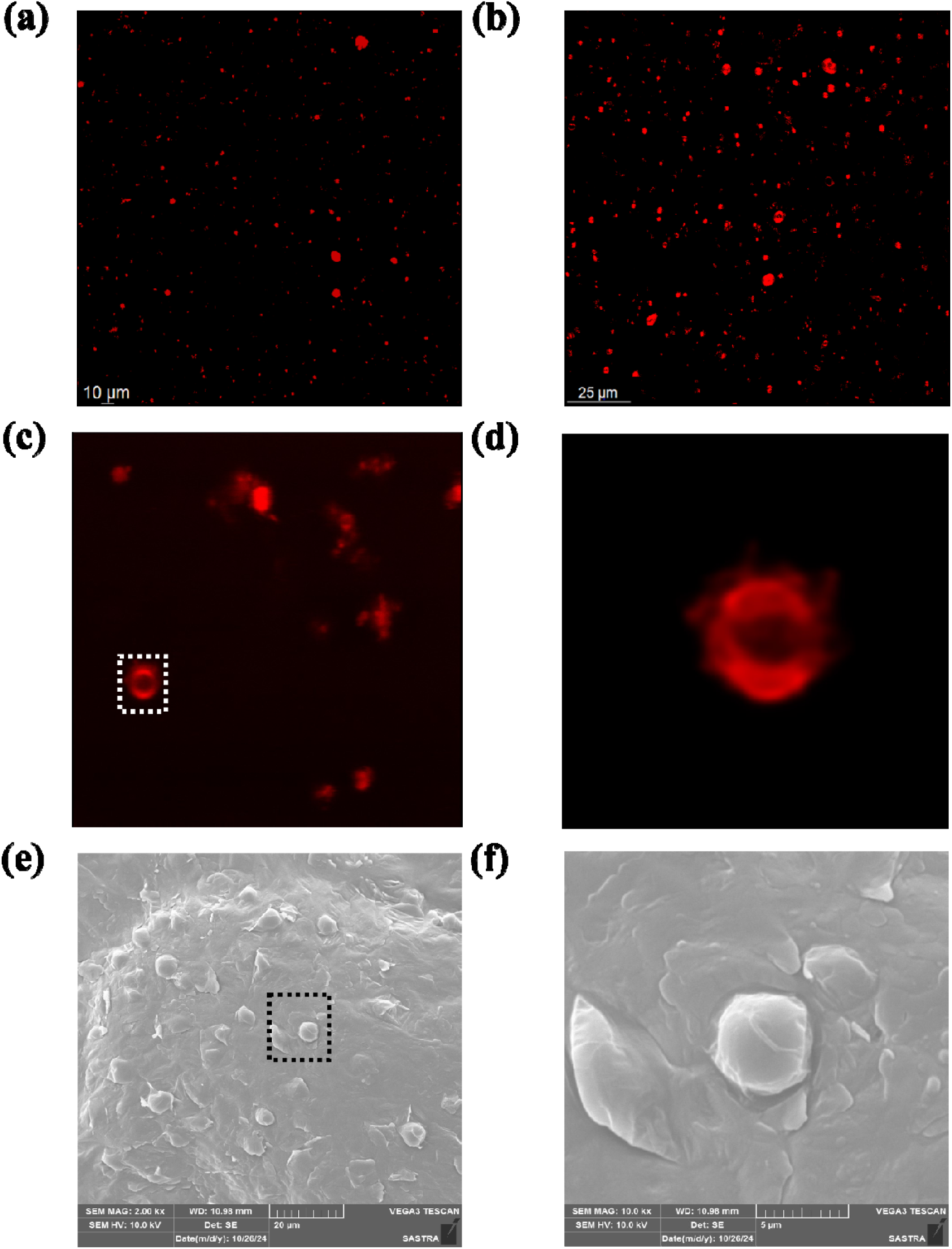
Confocal and SEM characterization of liposomes. (a,b) Confocal images at the 37th and 18th cycles of pulsed probe sonication (20% amplitude; 5 s ON / 55 s OFF). (c,d) Three-dimensional Z-stack and magnified view (dotted square) revealing lamellar structure indicative of the MLV phase. (e) SEM image at the 37th cycle. (f) Magnified view (black dotted square) showing spherical ULVs with slight membrane undulation.

These findings show that sonication amplitude and duty cycle govern vesicle morphology: high amplitude yields rapid ULV formation but risks thermal damage, whereas low-amplitude pulsed sonication achieves the same outcome more safely over longer times. Therefore, tuning sonication parameters enables controlled and reliable production of the desired vesicle type (S10 in SI).

## 4. Conclusion

A systematic and optimized workflow for liposome preparation is established, integrating thin-film formation, controlled drying, calibrated rehydration, vesicle downsizing, and advanced image analysis (Fig. S8 in SI). Container geometry markedly influences film uniformity and vesicle morphology: sharp-edged Eppendorf tubes produce heterogeneous films and broad vesicle distributions, whereas curved watch glasses yield comparatively uniform films; the most homogeneous and completely dried films are obtained by rotary evaporation at 41 °C and 480 mbar for 15 min (60–70 rpm), followed by 36 °C and 56 mbar for 10 min (70 rpm) and overnight desiccation at 866.6 mbar, while air drying often results in sticky, incompletely dried films. Rehydration is optimized at 4 mL HEPES buffer per 10 mg lipid, minimizing vesicle overlap while maintaining sufficient concentration; lower volumes increase overlap, whereas higher volumes reduce vesicle density. For MVV downsizing, bath sonication (up to 90 min) predominantly yields MLVs without contamination but requires prolonged processing. In contrast, probe sonication enables rapid size reduction; continuous operation at 30% amplitude for 30 s induces heating and metallic contamination. Pulsed sonication at 30% amplitude (10 s ON/50 s OFF, 9 cycles) reduces contamination but retains thermal effects, whereas further optimization to 20% amplitude (5 s ON/55 s OFF) for 37 cycles, corresponding to net ON times of 90 s and 185 s, reproducibly generates MLVs and ULVs, respectively, without detectable heating or probe-derived particles; residual titanium fragments are removed by centrifugation at 10,000 × g for 3 min. For SEM preparation, removal of excess buffer components, including sucrose, is essential to prevent charging during gold sputtering. Quantitative robustness is enhanced using a machine learning– based Python image analysis workflow. Collectively, the optimized protocol provides a reproducible, contamination-minimized, and thermally controlled strategy for precise regulation of liposome phase behavior, vesicle size distribution, and structural integrity.

## 6. Acknowledgements

A.P. acknowledges the support under Anusandhan National Research Foundation (ANRF), Department of Science and Technology, Government of India [SERB-SRG/2022/001489] and T.R. Rajagopalan research fund, SASTRA University, India. A.R., S.S. and A.P. would like to thank to Rajan K.S., Rajesh Y.B.R.D., Dhakshinamoorthy S., Shanker Jha, Senthilkumar R., Arunachalam J., Akshaya J. and their lab members for their support and valuable assistance.

## References

1. Adler K. and Schiemann J. (1985). Characterization of liposomes by scanning electron microscopy and the freeze-fracture technique. Micron and Microscopica Acta. 16(2):109–113. 10.1016/0739-6260(85)90039-5.

2. Andra V.V.S.N.L., Pammi S.V.N., Bhatraju L.V.K.P. and Ruddaraju L.K. (2022). A Comprehensive Review on Novel Liposomal Methodologies, Commercial Formulations, Clinical Trials and Patents. BioNanoScience. 12(1):274–291.10.1007/s12668-022-00941-x.

3. Aranda-Lara L., Morales-Avila E., Luna-Gutiérrez M.A., Olivé-Alvarez E. and Isaac-Olivé K. (2020). Radiolabeled liposomes and lipoproteins as lipidic nanoparticles for imaging and therapy. Chemistry and Physics of Lipids. 230:104934.10.1016/j.chemphyslip.2020.104934.

4. BanghamA. D., Hill M.W., and Miller N.G.A (1974). Preparation and Use of Liposomes as Models of Biological Membranes. In:Methods in membrane biology(Ed. Korn, E.D.). Springer, Boston, MA, 68pp.

5. Bibi S., Kaur R., Henriksen-Lacey M., McNeil S.E., Wilkhu J., Lattmann E., Christensen D., Mohammed A.R. and Perrie Y. (2011). Microscopy imaging of liposomes: From coverslips to environmental SEM. International Journal of Pharmaceutics. 417(1-2);138–150. 10.1016/j.ijpharm.2010.12.021.

6. Choi S., Kang B., Yang E., Kim K., Kwak M.K., Chang P.-S. and Jung H.-S. (2023). Precise control of liposome size using characteristic time depends on solvent type and membrane properties. Sci Rep. 13(1): 4728. 10.1038/s41598-023-31895-z.

7. Danaei M., Dehghankhold M., Ataei S., Hasanzadeh Davarani F., Javanmard R., Dokhani A., Khorasani S. and Mozafari M.R. (2018). Impact of Particle Size and Polydispersity Index on the Clinical Applications of Lipidic Nanocarrier Systems. Pharmaceutics. 10(2): 57. 10.3390/pharmaceutics10020057.

8. de Freitas C.F., Calori I.R., Tessaro A.L., Caetano W. and Hioka N. (2019). Rapid formation of Small Unilamellar Vesicles (SUV) through low-frequency sonication: An innovative approach. Colloids and Surfaces. B, Biointerfaces. 181:837–844.10.1016/j.colsurfb.2019.06.027.

9. Drabik D., Chodaczek G., Kraszewski S. and Langner M. (2020). Mechanical Properties Determination of DMPC, DPPC, DSPC, and HSPC Solid-Ordered Bilayers. Langmuir. 36(14): 3826–3835. 10.1021/acs.langmuir.0c00475.

10. Fulton M.D. and Najahi-Missaoui W. (2023). Liposomes in Cancer Therapy: How Did We Start and Where Are We Now. International Journal of Molecular Sciences. 24(7): 6615. 10.3390/ijms24076615.

11. Giuliano C.B., Cvjetan N., Ayache J. and Walde P. (2021). Multivesicular Vesicles: Preparation and Applications. ChemSystemsChem. 3(2): e2000049. 10.1002/syst.202000049.

12. Gomez A.G. and Hosseinidoust Z. (2020). Liposomes for Antibiotic Encapsulation and Delivery. ACS Infect Dis. 6(5): 896–908. 10.1021/acsinfecdis.9b00357.

13. Hadian Z., Sahari M.A., Moghimi H.R. and Barzegar M. (2014). Formulation, characterization and optimization of liposomes containing eicosapentaenoic and docosahexaenoic acids;a methodology approach. Iranian journal of pharmaceutical research. 13(2):393–404.PMID: 25237335

14. Has C. and Sunthar P. (2019). A comprehensive review on recent preparation techniques of liposomes. Journal of Liposome Research. 30(4):336–365. 10.1080/08982104.2019.1668010.

15. Kim E.-M. and Jeong H.-J. (2021). Liposomes: Biomedical Applications. Chonnam Med J. 57(1):27–35.10.4068/cmj.2021.57.1.27.

16. Liu P., Chen G., and Zhang J. (2022). A Review of Liposomes as a Drug Delivery System: Current Status of Approved Products, Regulatory Environments, and Future Perspectives. Molecules. 27(4): 1372. 10.3390/molecules27041372.

17. Lombardo D. and Kiselev M.A. (2022). Methods of Liposomes Preparation: Formation and Control Factors of Versatile Nanocarriers for Biomedical and Nanomedicine Application. Pharmaceutics. 14(3): 543.10.3390/pharmaceutics14030543.

18. Lu W.-L. and Qi X.-R. (2021). Liposome-Based Drug Delivery Systems. Springer Berlin Heidelberg. 10.1007/978-3-662-49320-5.

19. Lujan H., Griffin W.C., Taube J.H. and Sayes C.M. (2019). Synthesis and characterization of nanometer-sized liposomes for encapsulation and microRNA transfer to breast cancer cells. International Journal of Nanomedicine. 14:5159–5173.10.2147/IJN.S203330.

20. Maulucci G., De Spirito M., Arcovito G., Boffi F., Congiu Castellano A. and Briganti G. (2005). Particle Size Distribution in DMPC Vesicles Solutions Undergoing Different Sonication Times. Biophysical Journal. 88(5): 3545–3550. 10.1529/biophysj.104.048876.

21. Mehta M., Bui T.A., Yang X., Aksoy Y., Goldys E.M. and Deng W. (2023). Lipid-Based Nanoparticles for Drug/Gene Delivery: An Overview of the Production Techniques and Difficulties Encountered in Their Industrial Development. ACS Mater Au.3(6): 600–619. 10.1021/acsmaterialsau.3c00032.

22. Mendez R. and Banerjee S. (2017). Sonication-Based Basic Protocol for Liposome Synthesis. Methods in Molecular Biology. 1609:255–260.10.1007/978-1-4939-6996-8_21.

23. Morrissey J.H. (2001).Morrissey lab protocol for preparing phospholipid vesicles (SUV) by sonication. Urbana, IL: University of Illinois at Urbana-Champaign.

24. Naddaf Dezfuli S., Huan Z., Molc J.M.C., Leeflang M.A., Chang J. and Zhou J. (2014). Influence of HEPES buffer on the local pH and formation of surface layer during in vitro degradation tests of magnesium in DMEM. Progress in Natural Science. 24(5). 10.1016/j.pnsc.2014.08.009.

25. Nagayasu A., Uchiyama K. and Kiwada H. (1999). The size of liposomes: a factor which affects their targeting efficiency to tumors and therapeutic activity of liposomal antitumor drugs. Advanced Drug Delivery Reviews. 40(1): 75–87. 10.1016/S0169-409X(99)00041-1.

26. Nsairat H., AlShaer W.,Odeh F.,Essawi E., Khater D.,AlBawab A., El-Tanani M., Awidi A. and Mubarak M.S. (2023). Recent advances in using liposomes for delivery of nucleic acid-based therapeutics. OpenNano. 11: 100132. 10.1016/j.onano.2023.100132.

27. Robson A.-L., Dastoor P.C., Flynn J., Palmer W., Martin A., Smith D.W., Woldu A. and Hua S. (2018). Advantages and Limitations of Current Imaging Techniques for Characterizing Liposome Morphology. Frontiers in Pharmacology. 9:80. 10.3389/fphar.2018.00080.

28. Sessa G. and Weissmann G. (1968). Phospholipid spherules (liposomes) as a model for biological membranes. Journal of Lipid Research. 9(3): 310–318. PMID: 5646182.

29. Sharma K., Nilsuwan K., Ma L. and Benjakul S. (2023). Effect of Liposomal Encapsulation and Ultrasonication on Debittering of Protein Hydrolysate and Plastein from Salmon Frame. Foods. 12(4): 761. 10.3390/foods12040761.

30. Shohda K., Takahashi K. and Suyama A. (2015). A method of gentle hydration to prepare oil-free giant unilamellar vesicles that can confine enzymatic reactions. Biochemistry and Biophysics Reports. 3:76–82. 10.1016/j.bbrep.2015.07.005.

31. Šturm L. and PoklarUlrih N. (2021). Basic Methods for Preparation of Liposomes and Studying Their Interactions with Different Compounds, with the Emphasis on Polyphenols. International Journal of Molecular Sciences. 22(12): 6547. 10.3390/ijms22126547.

32. Umbarkar M., Thakare S., Surushe T., Giri A. and Chopade V. (2021). Formulation and Evaluation of Liposome by Thin Film Hydration Method. Journal of Drug Delivery and Therapeutics. 11(1): 72–76. 10.22270/jddt.v11i1.4677.

33. van der Veen J.N., Kennelly J.P., Wan S., Vance J.E., Vance D.E. and Jacobs R.L. (2017). The critical role of phosphatidylcholine and phosphatidylethanolamine metabolism in health and disease. Biochimica et Biophysica Acta (BBA) –Biomembranes. 1859(9): 1558–1572. 10.1016/j.bbamem.2017.04.006.

34. Wang N., Chen M. and Wang T. (2019). Liposomes used as a vaccine adjuvant-delivery system: From basics to clinical immunization. Journal of Controlled Release. 303: 130–150. 10.1016/j.jconrel.2019.04.025.

35. Zeng H., Guo S., Ren X., Wu Z., Liu S. and Yao X. (2023). Current Strategies for Exosome Cargo Loading and Targeting Delivery. Cells. 12(10):1416.10.3390/cells12101416.

36. Zhang G. and Sun J. (2021). Lipid in Chips: A Brief Review of Liposomes Formation by Microfluidics. International Journal of Nanomedicine. 16: 7391–7416. 10.2147/ijn.s331639.

37. Zhu T.F., Budin I. and Szostak J.W. (2013). Preparation of Fatty Acid or Phospholipid Vesicles by Thin-film Rehydration. In:Methods in Enzymology, Academic Press, 267–274.

