## Supplementary Information (SI) for "Sustainable Technology for the Fabrication of Liposomal Phases"

**S1. Microscopic techniques used for liposome characterization**

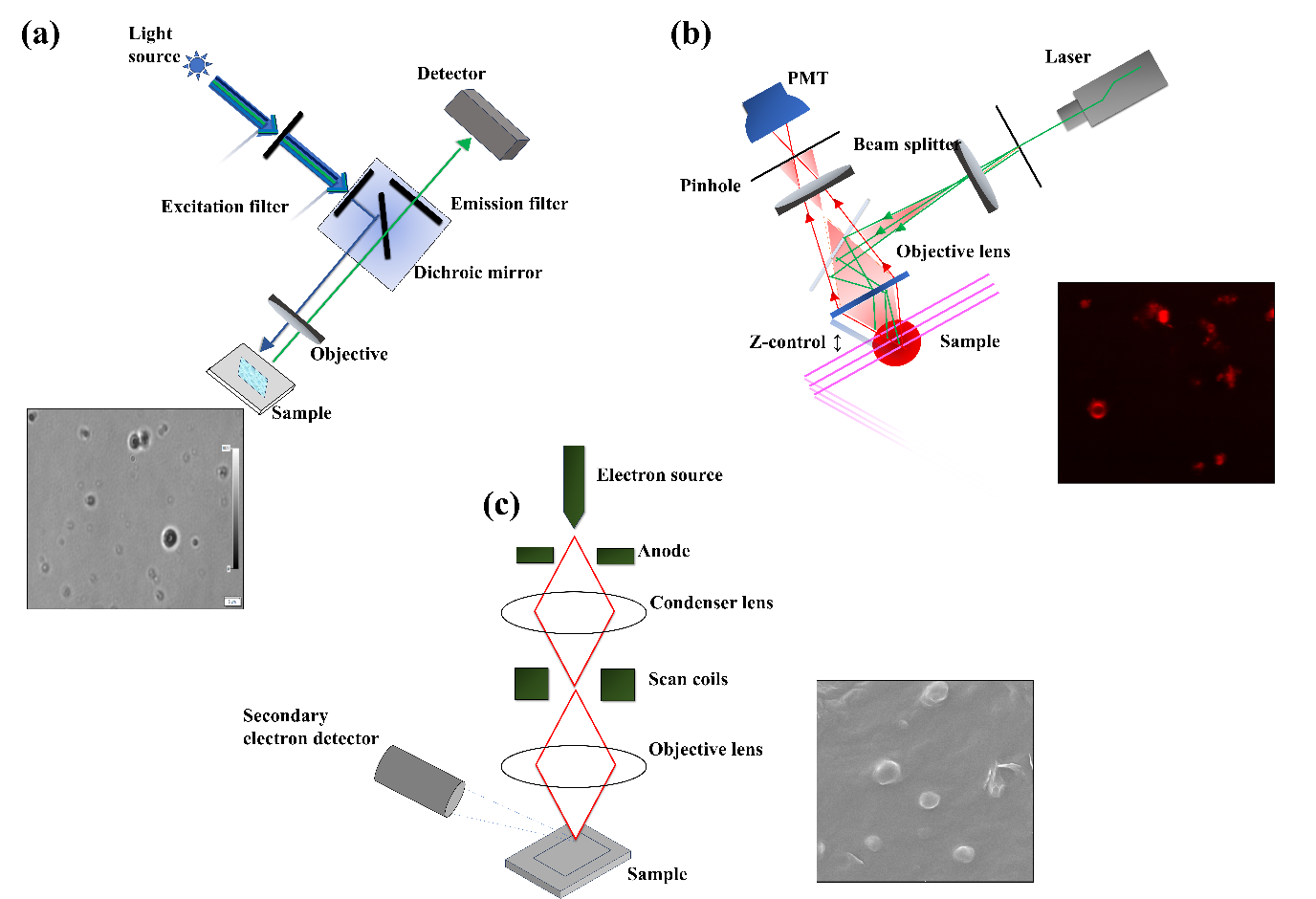

Figure S1: Schematic representation of the microscopy techniques used for liposome visualization: Bright-field microscopy (BF) imaging with an inverted fluorescence microscope shown in Fig. S1(a), used to determine vesicle size, shape, and population distribution; Confocal microscopy (CM) shown in Fig. S1(b), used to visualize internal structure and distinguish lamellarity of vesicles through z-stack imaging and Scanning electron microscopy (SEM) shown in Fig. S1(c), used to obtain high-resolution surface morphology of vesicles.

**S1.1. Direct visualization of liposomes using Bright-field Microscopy**

For Bright-field imaging, an IX73 Olympus inverted fluorescence microscopewas used. An aliquot of the sample (≈ 0.4µl) is placed on a glass slide, covered with a coverslip, and the slide is inverted such that the coverslip faces the objective lens. The sample is focused at 40X magnification, and images are captured using the Olympus DP23M monochrome digital camera and saved via cellSens software for subsequent analysis.

**Note:** To prevent the contamination, the cover slips are properly cleaned before use. For the inverted fluorescent microscopy, the slide should be placed inverted with the coverslip facing the objective lens.

**Disadvantage:** The main disadvantage of the direct visualization is that the samples dry quickly (Bibi *et al*., 2011), necessitating to capture the images immediately after placing the sample on the slide.

**S1.2. Visualization of Vesicles Using CM Studies**

The images of the liposomes are captured using the Stellaris 5 confocal microscope. The high-resolution images provide detailed visualizations of the morphology and size of the vesicle structures. Following a similar sample preparation method that is used for bright field microscopy, the vesicle sample is placed on a glass slide, covered with a coverslip, and observed under 40X objective magnification. Three-dimensional Z-stacks are recorded for liposomes and check the lamellarity of vesicles (as shown in Fig. 6 in main-text).

**Note:** To minimize the movement artifacts during the visualization of the liposomes using the laser confocal microscopy, it is recommended to use a confocal dish instead of slide and coverslip. The confocal dish provides a more stable imaging platform, reducing the potential movement of the vesicles and ensuring clearer and accurate imaging.

**S1.3. Visualization of ULV Using SEM Studies**

The electron microscopy (EM) techniques such as transmission electron microscopy (TEM) and scanning electron microscopy (SEM), are commonly used to study the liposomes, providing detailed information about their size, morphology (Robson *et al*., 2018), bilayer thickness, and inter-bilayer distance (Aranda-Lara *et al*., 2020; Andra *et al*., 2022).

The preparation of the sample for SEM study is carried out in several steps. A three-step centrifugation process was conducted to remove residual buffer components, followed by sputter coating and subsequent SEM visualization.

i. **Buffer Removal:** An ULV sample is centrifuged at 10,000 rpm for 30 minutes to pellet the vesicles. The supernatant containing the residual buffer is discarded using a micropipette.
ii. **Washing to remove excess buffer components**: The pellet is resuspended in the deionized water and the sample is centrifuged again at 10,000 rpm for 30 minutes and the supernatant is discarded to wash away remaining buffer components, such as sucrose or other solutes like NaCl, which could interfere with the SEM imaging. If the sucrose and other buffer contents are remained in the film, it can lead to the sample sticky and difficult to dry. This washing step is repeated twice.
iii. **Final washing step:** In the final wash, the sample undergoes centrifugation at 10,000 rpm for 15 minutes to ensure complete removal of buffer components while preserving the structural integrity of the vesicles.
iv. **Drying of film:** Once the washing process is completed, an aliquot of the sample (approximately 10µL), is placed on a 1x1 cm glass coverslip. The coverslip is then placed in a vacuum desiccator subjected to a pressure of 650mm Hg overnight. This drying step ensures complete dehydration of the sample, which is crucial to prevent any residual moisture, interfering with the SEM imaging.
v. **Sputter coating of film:** After drying, the sample is sputter-coated with a 4 nm thin layer of gold. This coating enhances conductivity, minimizing charging effects typically observed in the non-conductive biological samples, and improves image resolution and contrast.
vi. **Visualization of sample:** The dried sputter coated film is mounted on an SEM stub using a carbon tape and examined under SEM. The SEM allows for a detailed observation of the vesicles, including their size and morphology (Adler and Schiemann, 1985; Bibi *et al*., 2011; Robson *et al*., 2018; Lujan *et al*., 2019)

**S2. Protocol for Cleaning Glass Vials with Safety Considerations**

**S2.1 Protocol to clean glass vials**

Chromic sulfuric acid (also known as cleaning solution) is added to water and the glassware is completely immersed in the solution and left overnight to remove any organic residue from it. Then, the glassware is washed using the soap solution, air dried, and stored.

**S2.2 Safety and Handling**

Chloroform is volatile, toxic, and carcinogenic; therefore, all procedures involving chloroform were conducted in a certified chemical fume hood with appropriate personal protective equipment (PPE). Direct skin contact and inhalation were avoided. Glass pipettes were used to prevent solvent-induced degradation of plastic materials.

Many fluorescent dyes are also potentially carcinogenic; thus, appropriate protective measures were implemented. When fluorescent probes were incorporated, round-bottom flasks were wrapped in aluminium foil to prevent photodegradation.

Lipids stored at −20 °C were allowed to equilibrate to room temperature before use. To minimize lipid oxidation—particularly in mixtures containing unsaturated fatty acids—air-drying procedures were conducted in low-humidity environments, and lipid stocks were stored under inert atmosphere. The round-bottom flask was optionally rinsed with methanol prior to lipid addition to remove potential contaminants.

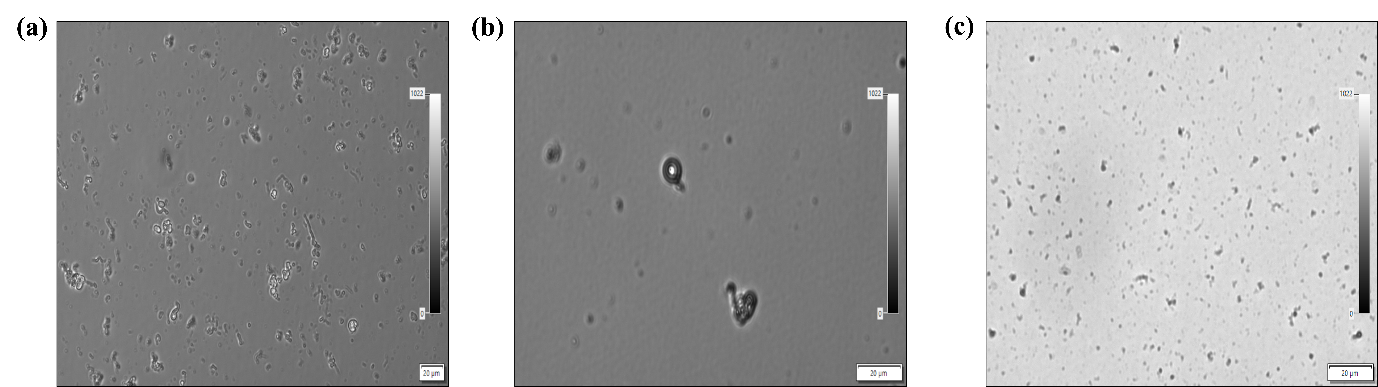

Figure S2: Bright-field images of lipid vesicles formed after hydrating films prepared byair-drying in an Eppendorf tube, exhibiting heterogeneous vesicle distribution shown in Fig. S2(a); while air-drying on a concave watch glass, exhibiting likely-uniform vesicle distribution shown in Fig. S2(b). Fig. S2(c) illustrates vesicles obtained by rotary evaporation, exhibiting a uniform and homogeneous distribution.

**S3. Thin film preparation by air dry method**

For the air-drying method, the lipid-chloroform solution is prepared in an Eppendorf tube and on the concave watch glass to investigate the influence of the container on film formation (Has and Sunthar, 2020). Both the Eppendorf tube with its lid open, and the watch glass are placed in the fume hood at room temperature for 7 to 8 hours to allow for the complete solvent evaporation. After chloroform evaporation, the DMPC lipid forms a film on the surfaces of Eppendorf tube and watch glass, respectively (Bangham *et al*., 1974). Upon rehydration of thin films from the air-drying method, the resulting lipid vesicles formed are visualized using bright-field microscopy.

**S4. Rehydration Buffer Preparation**

HEPES (N-2-hydroxyethylpiperazine-N-2-ethane sulfonic acid) buffer is used for rehydration of the thin film and contains 50 mM HEPES, 150 mM NaCl, and 5% (w/v) sucrose.

The required mass of each component for 20mL is calculated using:

Mass(g) = Concentration(mM) × Volume(L) ×Molar mass(g/mol)

HEPES=(50×10^−3^mol/L)×(20×10^−3^L)×(238.3g/mol)

=0.05 ×0.02×238.3g

=0.23g

NaCl = (150×10^−3^mol/L)×(20×10^−3^L)×(58.44g/mol)

=0.15 ×0.02×58.44g

=0.17g

Sucrose = 5%(w/v) ×20mL

=0.05g/mL×20mL

=1g

The component masses used for buffer preparation are summarized in Table 1 and are dissolved in 20mL of distilled water.

| Component | Amount (grams) |
| --- | --- |
| HEPES | 0.23 |
| NaCl | 0.17 |
| Sucrose | 1 |

Table 1: The buffer is prepared by dissolving the required components, in their respective masses, in 20 mL of distilled water.

The pH of the buffer solution can be adjusted using appropriate pH adjusting reagent, such as sodium hydroxide (NaOH) or hydrochloric acid (HCl). The reagent is added and mixed gradually and to adjust the pH until it reaches to 7.5.

**Caution:** It is important to handle with care, as HCl and NaOH are both highly irritating and corrosive.

**S5. Bath Sonication for MLV Formation**

For the bath sonication method, an Ultrasonic cleaner bath (Frequency: 40kHz) is employed. The bath sonicator is filled with the laboratory-grade distilled water. The Eppendorf tubes containing 0.5ml of vortexed suspension with MVVs are held upright in the water bath using a floater. The Eppendorf tubes containing the sample are fully immersed in the water bath to ensure efficient energy transfer during sonication (Morrissey, 2001). Then the bath sonication is carried out at the temperature of 37$℃$ for varying durations of 10, 20, 30, 45, 60, and 90 minutes (Hadian *et al*., 2014). At each sonication interval, the samples are periodically visualized under a microscope to monitor the formation of dis tinct phases of liposomes such as MVVs, MLVs and ULVs. The microscopic analysis reveals that the bath sonication up to 90 minutes predominantly results in MLVs along with residual MVVs, with no ULVs observed even at the longest time durations (Fig. S1).

**Advantage:** The bath sonication efficiently sonicates the liposome without contaminating the sample.

**Disadvantage:** A significant drawback is its longer processing time compared to the probe sonication method.

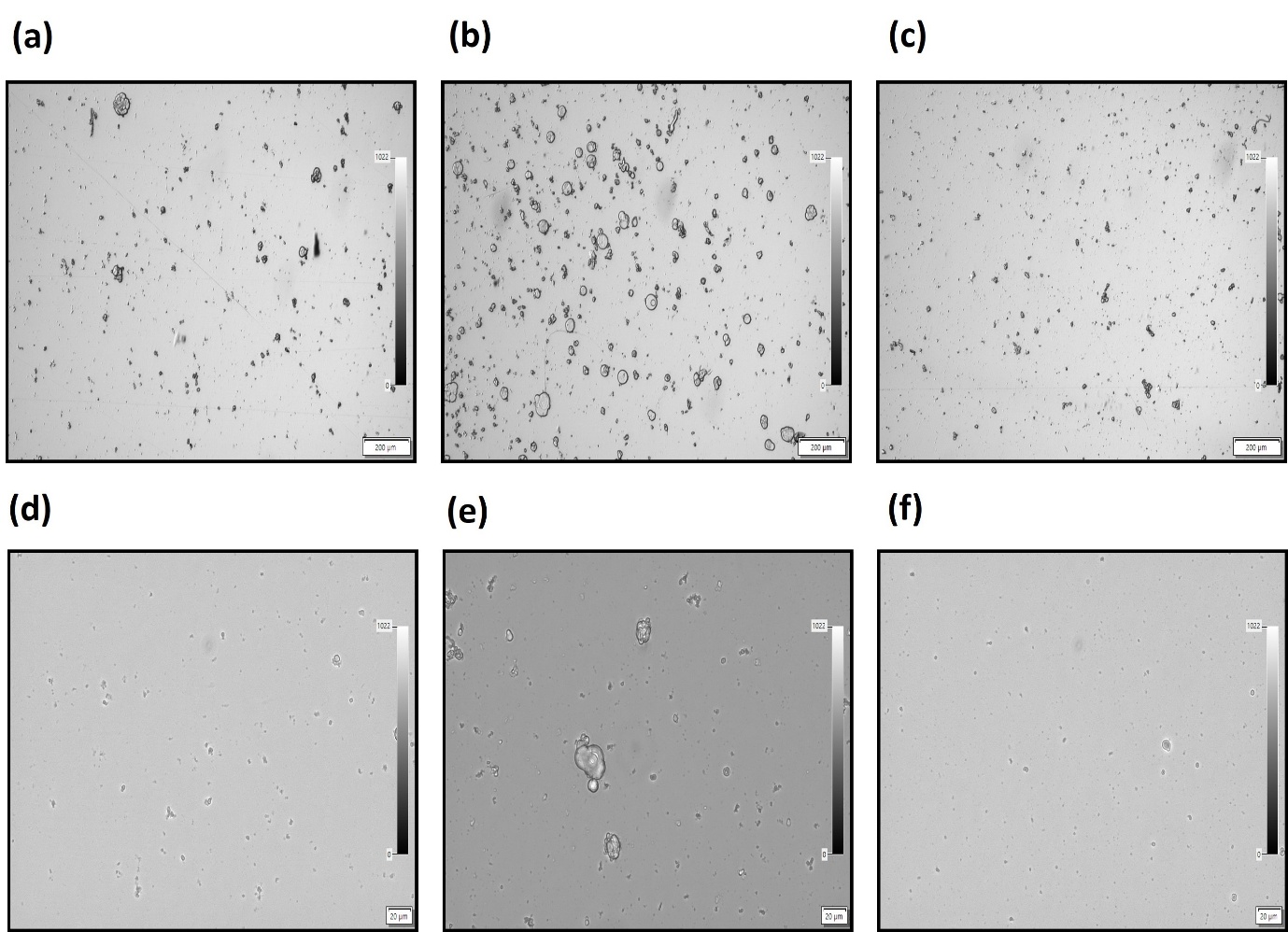

Figure S1: Vesicles under the bath sonication of varying time durations. The bath sonication of MVVs (after vortexing) is carried out which exhibits the downsizing of the vesicles at different time intervals: 10 minutes (Fig. S1(a)), 20 minutes (Fig. S1(b)), 30 minutes (Fig. S1(c)), 45 minutes (Fig. S1(d)), 60 minutes (Fig. S1(e)), 90 minutes (Fig. S1(f)), respectively.

**S6. Python-based image analysis for vesicle size estimation**

A Python-based image analysis workflow, utilizing OpenCV, a machine learning software library, is employed to analyze the image of the vesicles and study the size and the morphology of the vesicles. The images are pre-processed by applying the Gaussian blur filter to reduce noise, followed by histogram equalization to enhance the contrast. The vesicle boundaries are detected using the Canny-Edge-Detection algorithm, and the contours are extracted. For each detected contour, the area and the radius are calculated. The pixel measurements are converted to micrometers using a predefined scale factor for the accurate vesicle size quantification. To investigate the distribution of the vesicle sizes, the probability distribution of the vesicle radius was calculated. To the histogram data, Gaussian fitting is applied, enabling us to track the mean vesicle size with errors.

**S7. Different phases of liposomes—MVV, MLV and ULV**

The preparation of different liposomal phases via rehydration of lipid thin films was investigated and subsequently characterized, as shown in Fig. S2.

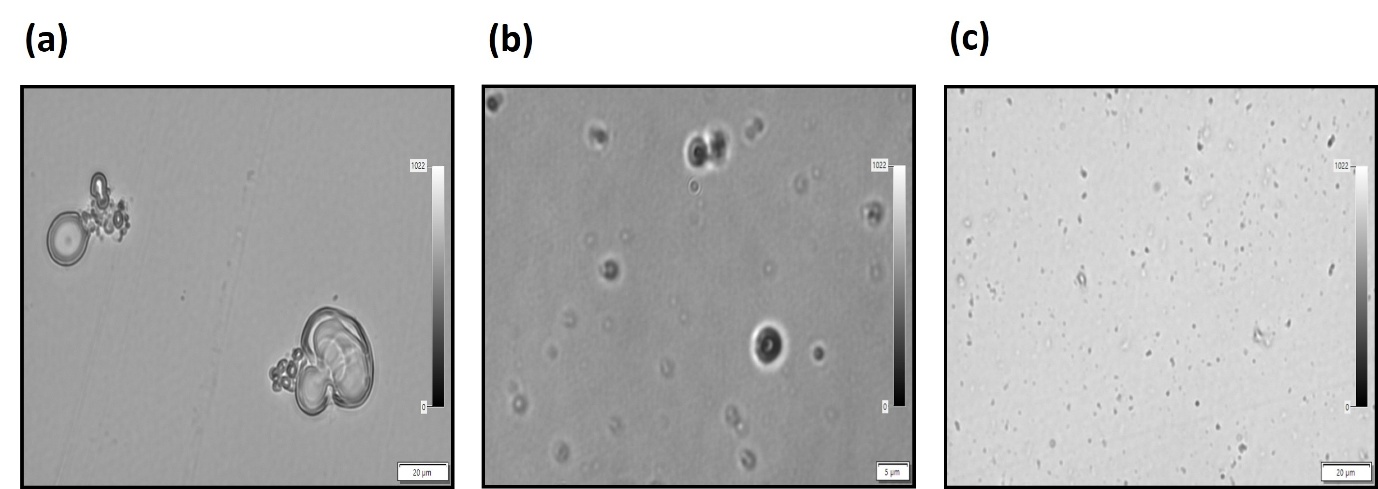

Figure S2: (a) shows that the MVVs are exhibited after vortexing the hydrated film. (b) shows that after vortexing, the MLVs are exhibited after 30 seconds of continuous probe sonication at 30% amplitude. (c) shows unilamellar vesicles formed after 185 seconds of pulsed probe sonication at 20% amplitude (5-second ON / 55-second OFF per cycle).

**S8. Area and radius of MVV, MLV and ULV**

Images of MVV, MLV, and ULV phases were collected and the probability distributions of vesicle area (A) and radius (R) were calculated, as presented in Fig. S3.

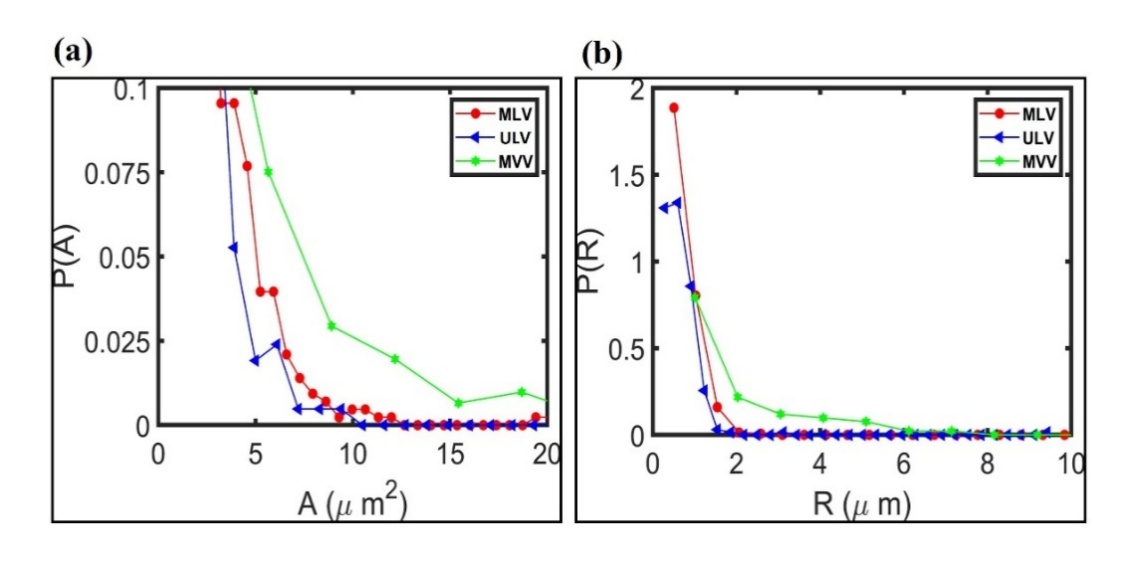

Figure S3: Fig. S3(a) and Fig. S3(b) show probability distribution of area, A and radius, R of MVV, MLV and ULV, respectively, showing the size of MVV > MLV >ULV

**S9. Optimization of Liposome Size via Pulsed Probe Sonication at Different Amplitudes**

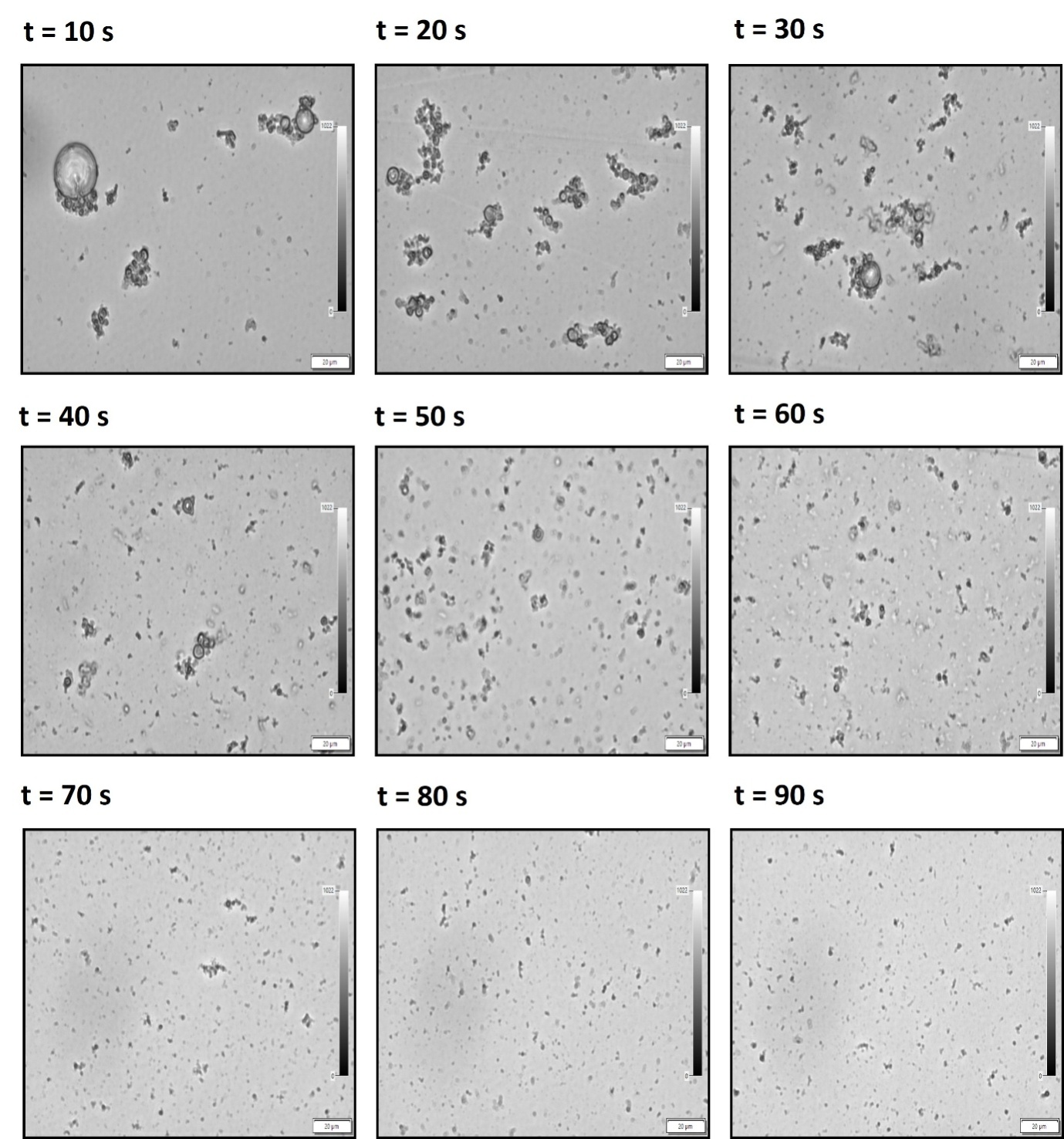

Figure S4: Vesicles under the pulsed probe sonication at 30% amplitude of varying ON time durations. The pulsed probe sonication of MVVs (after vortexing) is carried out, resulting in the downsizing of the vesicles at different ON time intervals, t. Fig. S4 shows large multilamellar vesicles (MLVs) present at early time points (t = 10 to 30 s) which gradually downsize into smaller unilamellar vesicles (ULVs). The ULVs begin to form after a net ON-time of 60 seconds. A clear reduction in size polydispersity and an increase in vesicle uniformity are observed at longer ON-times (t = 70 to 90 s)

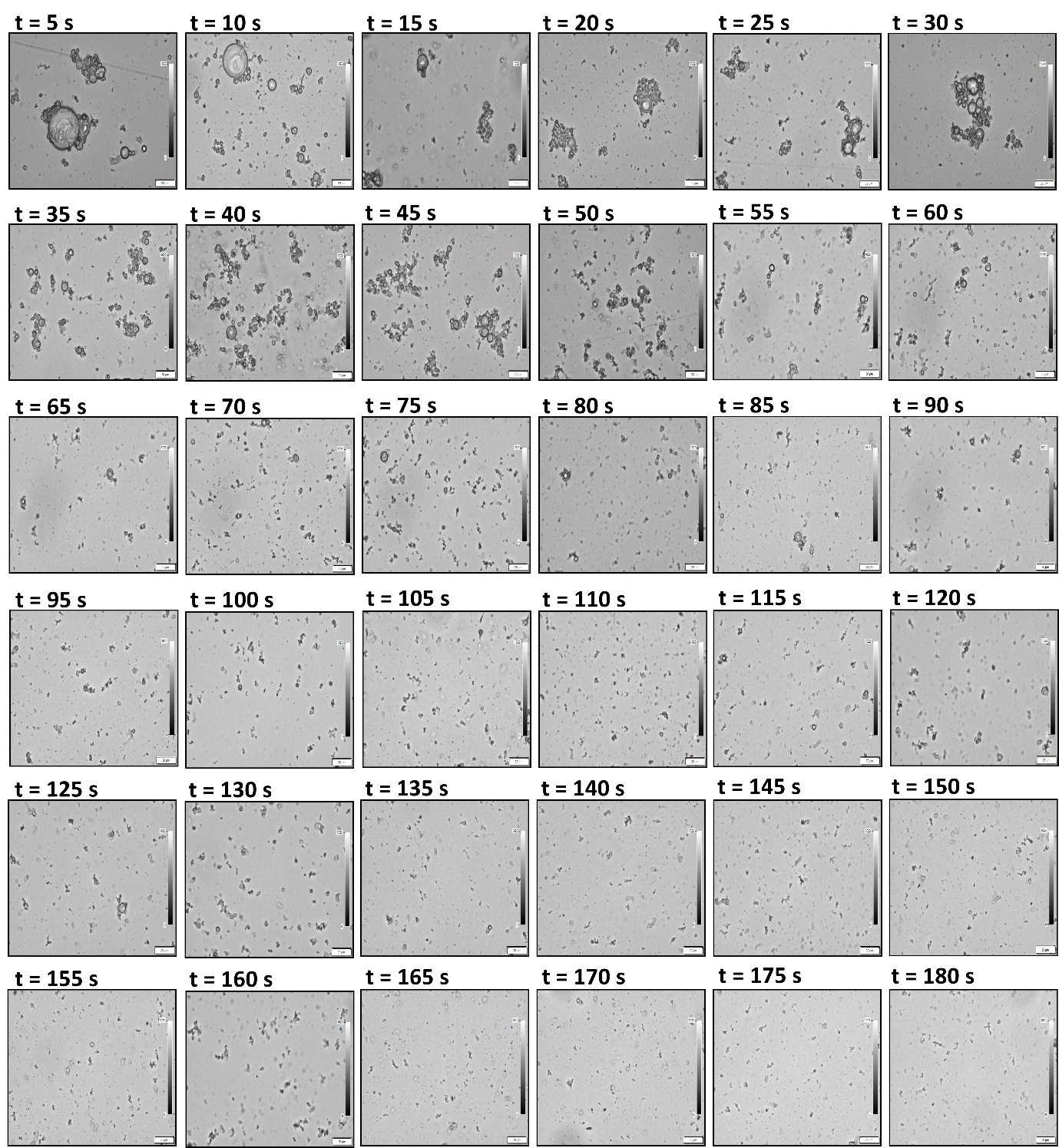

Figure S5: Vesicles under the pulsed probe sonication at 20% amplitude of varying ON timedurations. The pulsed probe sonication of MVVs (after vortexing) results in downsizing of the vesicles at different ON time intervals, t. A clear reduction in size polydispersity and an increase in vesicle uniformity are observed at a late time (t = 70 seconds). Fig. S5 reveals that large multilamellar vesicles (MLVs) begins to appear at early ON-time points (t ∼ 25 s) and MLVs are evident at 90 seconds (18^th^ cycle). The liposomes gradually downsize and the small unilamellar vesicles (ULVs) begin to be exhibited at 160 seconds (32nd cycle) and are evident at 185 seconds (37^th^ cycle). For every time point, 20 images are recorded and only one representative image from each time point is shown.

**S10. Experimental Considerations, Advantages, and Limitations**

**S10.1 Considerations During Optimization of Rehydration Buffer Volume**

During optimization of the rehydration buffer volume, rehydration was performed at temperatures above the main phase transition temperature to ensure efficient lipid swelling and vesicle formation (Lombardo and Kiselev, 2022). In addition, effective dissolution of the thin phospholipid film required the presence of metal ions in the buffer. Accordingly, NaCl-containing buffer was used, as ionic strength facilitated lipid hydration and dispersion (Zhu et al., 2013).

**S10.2 Considerations During Optimization of Probe Sonication for Different Liposomal Phases**

During optimization of probe sonication to obtain different liposomal phases, several technical considerations were addressed. Multilamellar vesicle (MLV) samples were stored at 4 °C for up to 6 weeks without significant degradation. Alternatively, unilamellar vesicles could be prepared using a membrane extruder, where vesicle size is defined by the membrane pore size (Andra et al., 2022); however, limitations of extrusion include difficulty in large-scale production and filter clogging. Following probe sonication, samples were centrifuged at 10,000 × g for 3 min to remove residual titanium particles from the probe tip and any unreconstituted lipids.

Probe sonication inherently generates heat, which may cause phospholipid degradation at the probe surface and potential metal leaching into the solution (Mendez and Banerjee, 2017); therefore, temperature was continuously monitored to prevent thermal effects. A primary advantage of probe sonication was the rapid production of unilamellar vesicles with reduced processing time. A limitation was the possible release of metallic particles into the liposome suspension; however, controlled pulsed operation and post-sonication centrifugation minimized both thermal effects and contamination risks.

**S11 Schematic Workflow for Controlled Generation of Distinct Liposomal Phases**

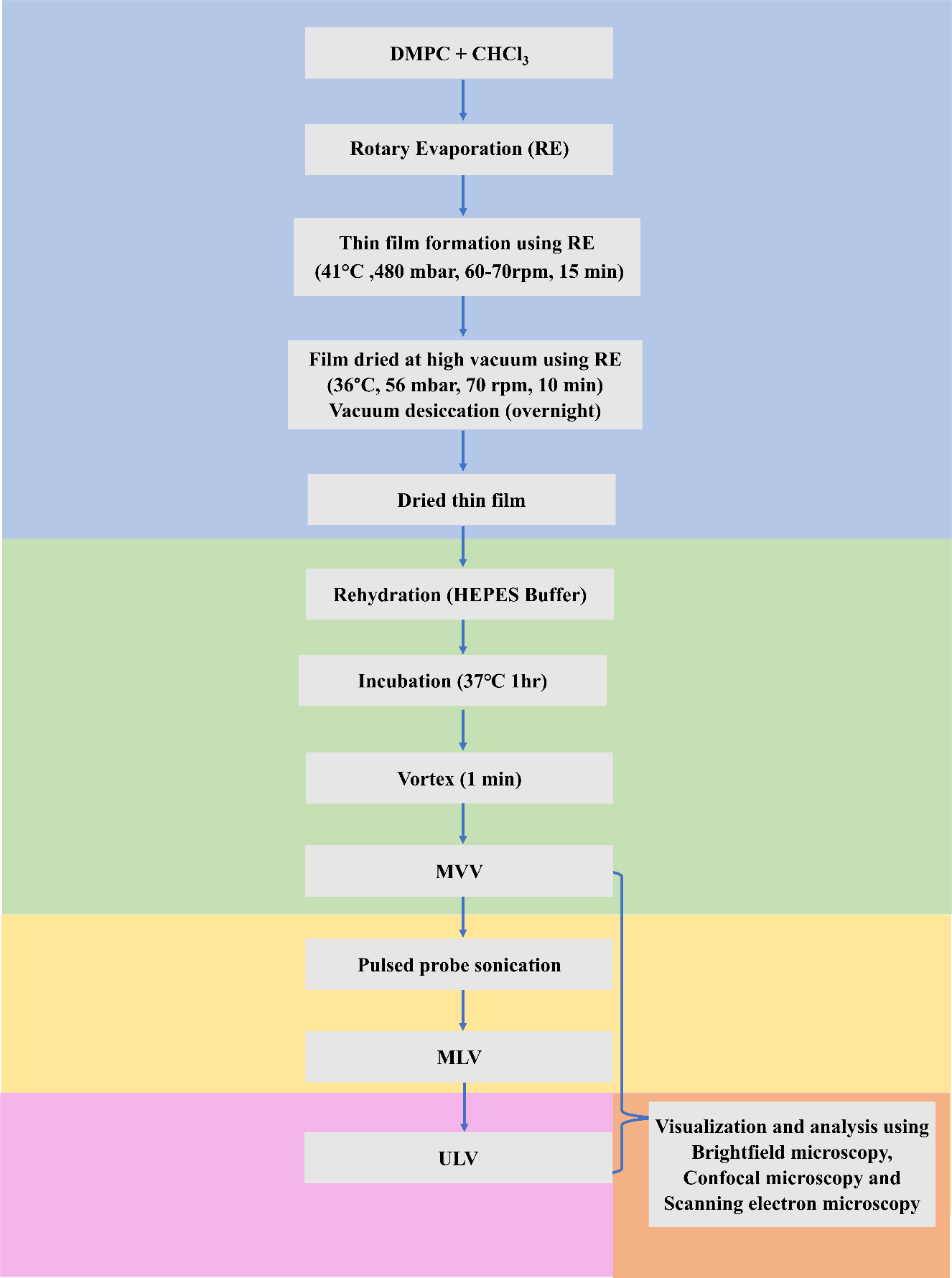

*Figure S8: Flowchart of the methodology for generating distinct phases of liposomes. The flowchart illustrates the sequential preparation of different phases of liposome, divided into five main steps highlighting with different colors. The blue, green, yellow, and pink represent thin-film formation by rotary evaporation technique, MVV formation, MLV formation, and ULV formation, respectively. The final step of visualization and characterization is depicted in orange.*
